# Novel Oral Adjuvant to Enhance Cytotoxic Memory-Like NK Cell Responses in an HIV Vaccine Platform

**DOI:** 10.1101/2024.05.11.593683

**Authors:** Mario Alles, Manuja Gunasena, Christina Isckarus, Ilmini De Silva, Sarah Board, Will Mulhern, Patrick L. Collins, Thorsten Demberg, Namal P.M. Liyanage

## Abstract

Antibody-dependent cell-mediated cytotoxicity, mediated by natural killer (NK) cells and antibodies, emerged as a secondary correlate of protection in the RV144 HIV vaccine clinical trial, the only vaccine thus far demonstrating some efficacy in human. Therefore, leveraging NK cells with enhanced cytotoxic effector responses may bolster vaccine induced protection against HIV. Here, we investigated the effect of orally administering indole-3-carbinol (I3C), an aryl hydrocarbon receptor (AHR) agonist, as an adjuvant to an RV144-like vaccine platform in a mouse model. We demonstrate the expansion of KLRG1-expressing NK cells induced by the vaccine together with I3C. This NK cell subset exhibited enhanced vaccine antigen-specific cytotoxic memory-like features. Our study underscores the potential of incorporating I3C as an oral adjuvant to HIV vaccine platforms to enhance antigen-specific (memory-like) cytotoxicity of NK cells against HIV-infected cells. This approach may contribute to enhancing the protective efficacy of HIV preventive vaccines against HIV acquisition.

## Introduction

Antibody-dependent cell-mediated cytotoxicity (ADCC) is a critical mechanism involving certain immune cells, particularly natural killer (NK) cells and monocytes, in destroying virally infected cells coated with specific antibodies. This process entails the interaction between the Fc portion of antibodies and Fc receptors (FcRs) expressed on the surface of these effector cells. ADCC has emerged as an important immune correlate in recent vaccine trials, including the RV144 trial, which demonstrated some degree of protection against HIV acquisition ^1, 2^. This study utilized a combination of a live-attenuated recombinant canarypox virus vector encoding HIV-1 clade E env and clade B gag and protease (ALVAC-HIV) along with AIDSVAX B/E gp120/alum, demonstrating a moderate efficacy of 31.2% over a 3.5-year period ^3^. We recapitulated the RV144 like study in rhesus macaque models and demonstrated similar vaccine efficacy in SIV infection^4–10^.

Natural killer (NK) cells, the primary effectors of ADCC in vivo, surveil the body for virally infected cells ^11^. Upon activation, NK cells release effector molecules such as perforins and granzyme B, leading to target cell death ^12^. Additionally, they can induce apoptosis in virally infected target cells through the FAS and tumor necrosis factor-related apoptosis inducing ligand (TRAIL) pathways ^13^. Exploration of unique NK cell phenotypes may offer novel insights into their variable cytotoxic abilities and potential for enhancing their antiviral effector mechanisms. For instance, Killer cell lectin-like receptor subfamily G member-1 (KLRG1) is a receptor expressed on NK cells and is also considered a marker of NK cell memory and maturity ^14^. KLRG1^+^ NK cells are also known to display greater NK cell effector responses such as ADCC, compared to their KLRG1^-^ counterparts. KLRG1^+^ NK cells exhibited higher cytotoxicity and IFN-γ release via antibody-dependent cellular cytotoxicity compared to KLRG1^−^ NK cells ^15^. Similarly, differential expression of other receptors of NK cell activity like CD27 and CD11b, along with their differential ability to degranulate on exposure to target HIV antigens may support their functional alterations that may be useful in mediating anti-HIV responses ^16^.

Investigations into enhancing clinically relevant NK cell effector responses to augment antibody-based immunotherapy are already underway within the field of cancer therapeutics ^17^. Similarly, harnessing NK cell-mediated antiviral mechanisms for HIV through vaccination could represent a pivotal advancement in developing efficacious anti-HIV vaccine strategies.

The aryl hydrocarbon receptor (AHR) is a ligand-activated transcription factor present in various immune cells including NK cells. Its activation controls the expression of several signaling molecules, particularly related to tumor immunology ^18^. However, less is known regarding its role in regulating effector functions of NK cells. Studies have shown that mice deficient in AHR have impaired anti-tumor responses in vivo, while mice treated with AHR agonists were more capable of controlling tumor growth ^19^. Given the importance of NK cells in cancer immunology and the role in promoting NK cytotoxicity through mechanisms like ADCC in anti-tumor activity, understanding the role of AHR agonists in enhancing NK effector responses may be useful for HIV vaccine development. AHR agonists may promote NK cytotoxicity against virally infected cells by increasing the effector functions of NK cells.

In this study, using a mouse model, we orally administered inodol-3-carbinol (I3C), a naturally occurring AHR agonist, as an adjuvant to an ALVAC-SIV/gp120-based (RV144-like) vaccine platform to observe its effect in modulating anti-HIV vaccine responses in the spleen. Here, we identified a subset of KLRG1-expressing NK cells that expanded in response to the vaccine following I3C co-administration, which exhibited enhanced vaccine antigen-specific cytotoxic memory-like features.

Overall, our study demonstrates that incorporating I3C as an oral adjuvant to an RV144-like vaccine platform may enhance anti-HIV/SIV NK cell cytotoxicity. This may potentially boost NK effector mechanisms, including ADCC, further increasing protection afforded by the vaccine against HIV acquisition. This novel concept may inspire the exploration of similar strategies for other HIV and infectious disease vaccine platforms.

## Results

### The Effects of the Oral Adjuvant I3C on splenic NK Cell Dynamics in ALVAC-SIV/gp120/alum-Immunized Mice

In this study, 12 C57BL/6 aged sex-matched mice were divided into two groups. All mice received an initial ALVAC-SIV vaccination at week 0, followed by a booster at week 2 with ALVAC-SIV and monomeric gp120 in alum. Additionally, six mice (group B) were administered oral I3C daily for four weeks beginning on week 0. After four weeks, all mice were euthanized, and splenic immune cells were analyzed using multi-dimensional flow cytometry to compare NK cells between the Vaccine + I3C and vaccine-only groups (Figure 1a). FlowSOM-based clustering and Uniform Manifold Approximation and Projection (UMAP) were employed for dimensional reduction visualization. Subsequently, CD45^+^CD3^-^CD19^-^NK1.1^+^ cells were analyzed, identifying 14 distinct NK cell clusters (Figure 1b). Expression intensity was correlated with clusters to delineate various NK cell subsets (Figure 1c). Statistical analyses revealed significant difference in the individual clusters between the two groups. Specifically, 10 (Cluster 2, 3, 4, 5, 6, 7, 10, 11, 12 and 13) out of the 14 clusters exhibited significant differences in expression (Figures 1d-g and Supplementary Figures 1b-k) between the two groups (p<0.05). Notably, two clusters (Cluster 5 and 13) in the Vaccine + I3C group showed significantly high expression of KLRG1, a marker associated with NK cell activation, maturation, and memory ^14,15,20^. These findings suggest that concurrent administration of I3C with the vaccine may enrich memory-like NK cell phenotypes with potentially enhanced activation and maturation levels.

**Figure 1.**
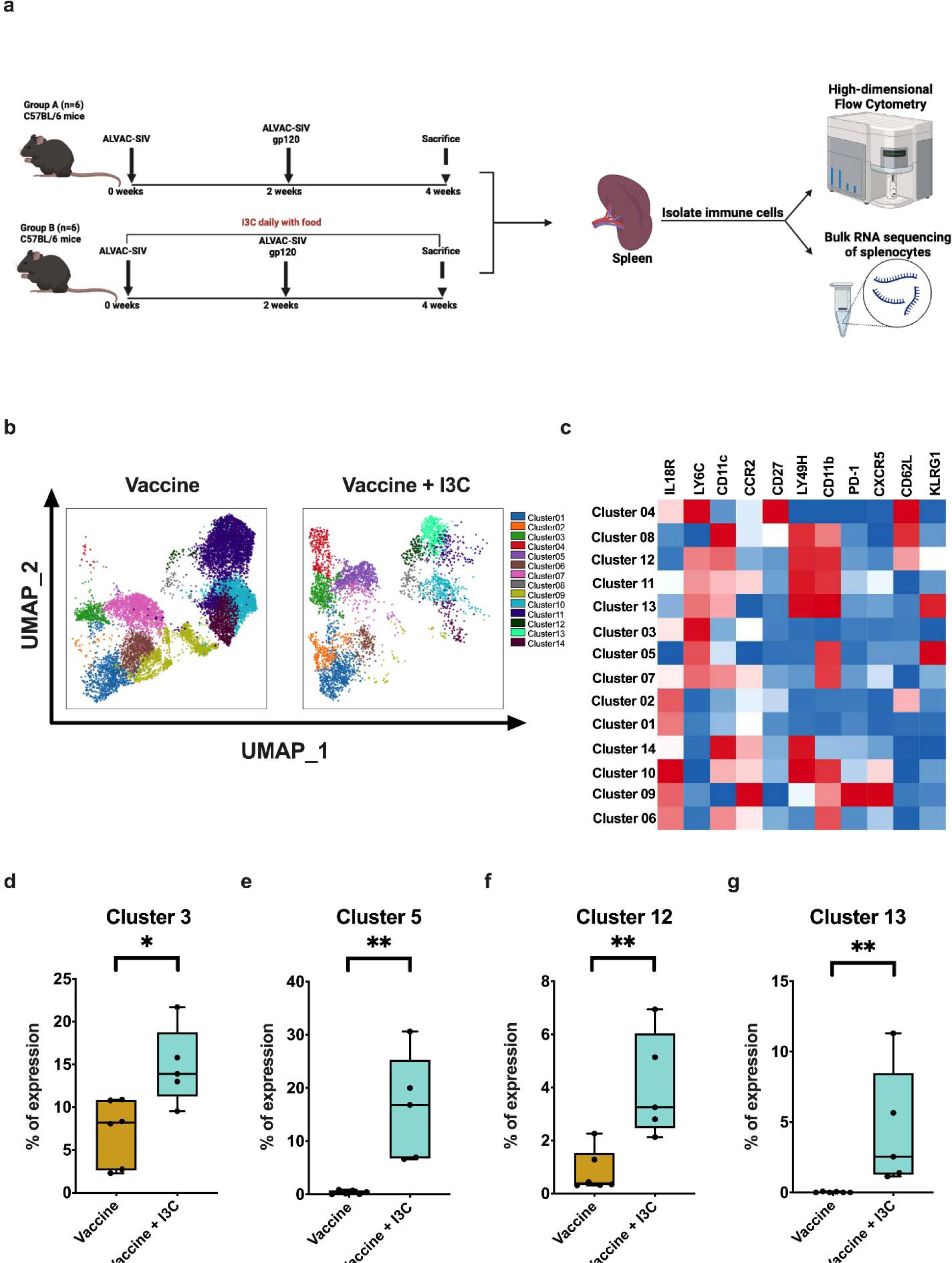
Differential characteristics of splenic NK cell populations between mice administered with an RV144-like vaccine regimen with oral adjuvant I3C (Vaccine + I3C group), and mice administered the vaccine only (Vaccine group). **(a)** Experimental design utilizing C57BL/6 mice. **(b)** FlowSOM-based cluster analysis of viable splenic CD45^+^CD3^-^ CD19^-^NK1.1^+^ cells of Vaccine and Vaccine + I3C groups depicted on a Uniform Manifold Approximation and Projection (UMAP). **(c)** Clusters-by-marker heatmap characterizing the receptor expression patterns of individual clusters. **(d-g)** Box plots depicting NK cell cluster frequency differences between Vaccine and Vaccine + I3C groups. The p-value was calculated using Wilcoxon rank-sum test; *p<0.05, **p<0.01. I3C, Indole-3-carbinol.

### Use of I3C as an oral adjuvant leads to the Expansion of KLRG1^+^NK Cells in Vaccinated Mice

We next employed conventional flow cytometry gating (Figure 2a and Supplementary Figure 2a) to analyze the differential expression of surface receptors in NK cells from both Vaccine and Vaccine + I3C groups. Our conventional gating analysis also unveiled a significant expansion of KLRG1^+^ NK cells within the Vaccine + I3C group when compared to the Vaccine group (p<0.01) (Figure 2b). To gain deeper insights into the characteristics of the expanded KLRG1^+^ NK cell subset and compare phenotypic features with KLRG1^-^ NK cells, we performed t-distributed stochastic embedding (t-SNE) dimensionality reduction analysis on CD45^+^CD3^-^ CD19^-^NK1.1^+^ cells (Figure 2c). This analysis enabled visualization of receptor expression pattern across KLRG1^+^ and KLRG1^-^ subsets, revealing distinct distribution and levels of surface markers such as Ly6C, CD27, CD11b, PD-1, CD11c, Ly49H, CXCR5, and CCR2 (Figures 2d-k). Specifically, KLRG1^+^ NK cells exhibited significantly higher expression of CD11b (p<0.01) and lower expression of CD27, indicating an enhanced cytotoxic potential ^16^. Additionally, their higher expression of CD11c (p<0.05) suggested an overall activated phenotype with heightened effector functions compared to CD11c^-^ cells ^21^. The higher expression of Ly6C (p<0.01) in the KLRG1^+^ NK subset suggests their potential to rapidly convert to Ly6C^low^ cells with stronger effector functions upon IL-15 stimulation ^22,23^. Given the association of Ly49H expression with NK cell memory, a higher expression of Ly49H (p<0.01) in the KLRG1^+^ NK subset would suggest a greater capacity to develop innate memory compared with KLRG1^-^ NK cells ^24^. This memory phenotype is further supported by upregulation of Ly6C and downregulation of CD27 (as mentioned previously) in KLRG1^+^ NK cells ^14^. Additionally, significant changes in the expression of chemokine receptors like CXCR5 and CCR2 between KLRG1^-^ and KLRG1^+^ NK cells (p<0.01), suggest differential chemotactic capabilities of these NK phenotypes^25^, further demonstrating that these expanded KLRG1^+^ NK cells are likely to be a phenotypically and functionally distinct NK cell subset.

**Figure 2.**
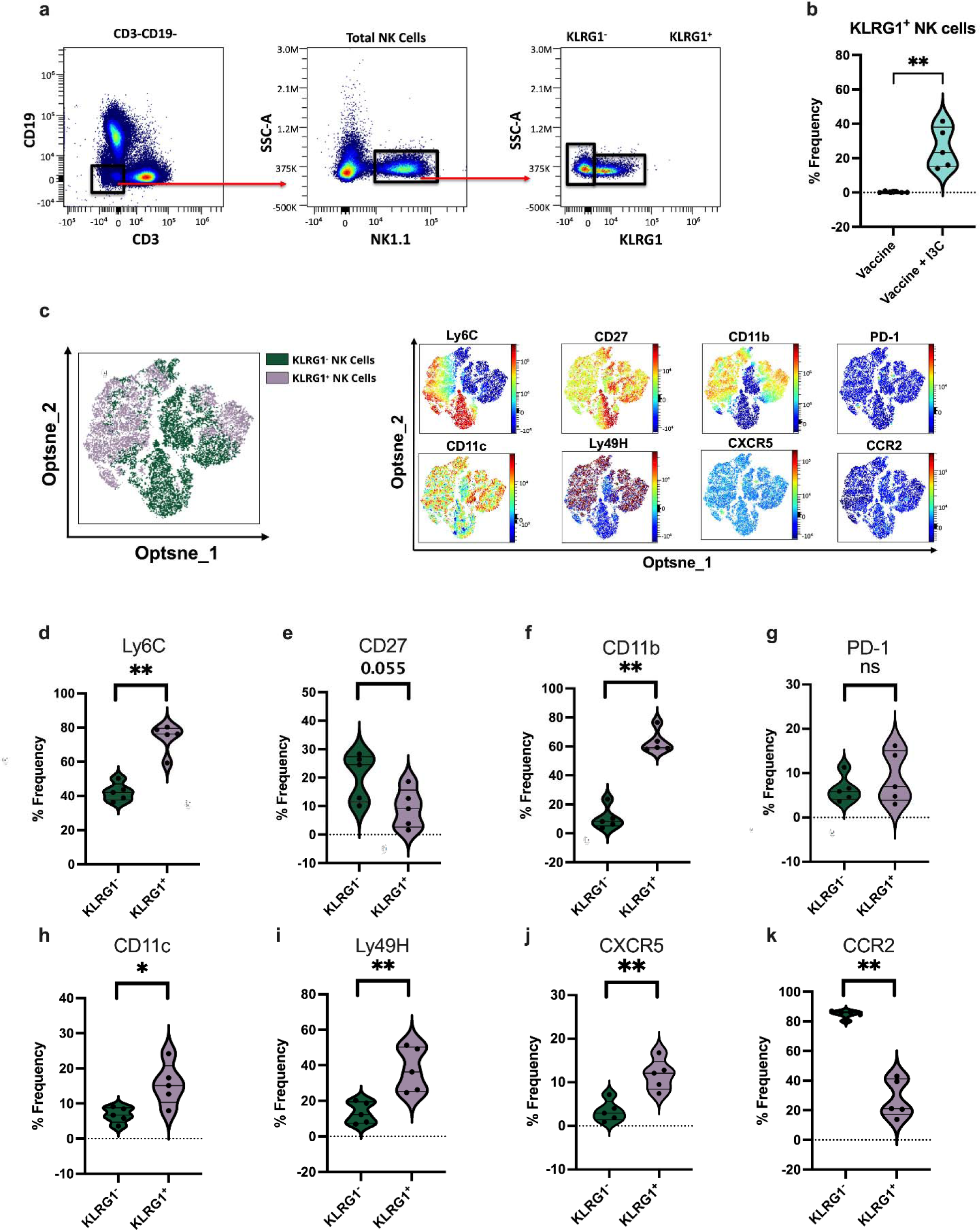
Phenotypic differences are observed between splenic KLRG1^-^ and KLRG1^+^ NK cells in Vaccine + oral adjuvant I3C administered mice. **(a)** Representative pseudo color plots depicting the gating strategy for KLRG1^+^ NK cells. **(b)** Violin plot illustrating differential proportions of KLRG1^+^ NK cells (among total NK cells) between Vaccine and Vaccine + I3C groups. **(c)** T-distributed stochastic neighbor embedding (t-SNE) analysis of concatenated samples gated on total viable NK1.1^+^ cells highlighting KLRG1^-^ and KLRG1^+^ NK cell subsets in the Vaccine + I3C group. Differential expression of Ly6C, CD27, CD11b, PD-1, CD11c, Ly49H, CXCR5 and CCR2 between KLRG1^-^ and KLRG1^+^ NK cell populations in the Vaccine + I3C group are visualized by t-SNE heatmaps. **(d-k)** Violin plots comparing frequencies of receptor expression between KLRG1^-^ and KLRG1^+^ NK cells in the Vaccine + I3C administered group. The p-value was calculated using Wilcoxon rank-sum test; *p<0.05, **p<0.01.

### Enhanced Cytotoxic Responses to SIV Peptides are Observed in KLRG1^+^ NK Cells in the SIV Vaccine + I3C Group

To determine the differential cytotoxic responses between KLRG1^-^ and KLRG1^+^ NK cells in the Vaccine + I3C group upon re-exposure to vaccine antigen, we conducted an in vitro experiment. Splenic mononuclear cells of these mice were stimulated with SIV-Gag peptides for 20 hours, as described in the materials and methods (Figure 3a). Additionally, we ran a parallel experiment exposing cells to phorbol 12-myristate 13-acetateand ionomycin (PMA/I) representing non-specific stimulation. Here, our results indicate that upon exposure to SIV-Gag, KLRG1^+^ NK cells demonstrated a significantly greater increase in their degranulation capability, as evidenced by greater increases in CD107a expression, compared to KLRG1^-^ NK cells (p<0.01) (Figures 3b and c). However, we did not see a significant difference in CD107a expression between KLRG1^-^ and KLRG1^+^ NK subsets when stimulated with PMA/I (Figure 3c), suggesting that the enhanced degranulation response may be a recall response of KLRG1^+^ NK cells to vaccine antigens signifying development of innate immune memory.

**Figure 3.**
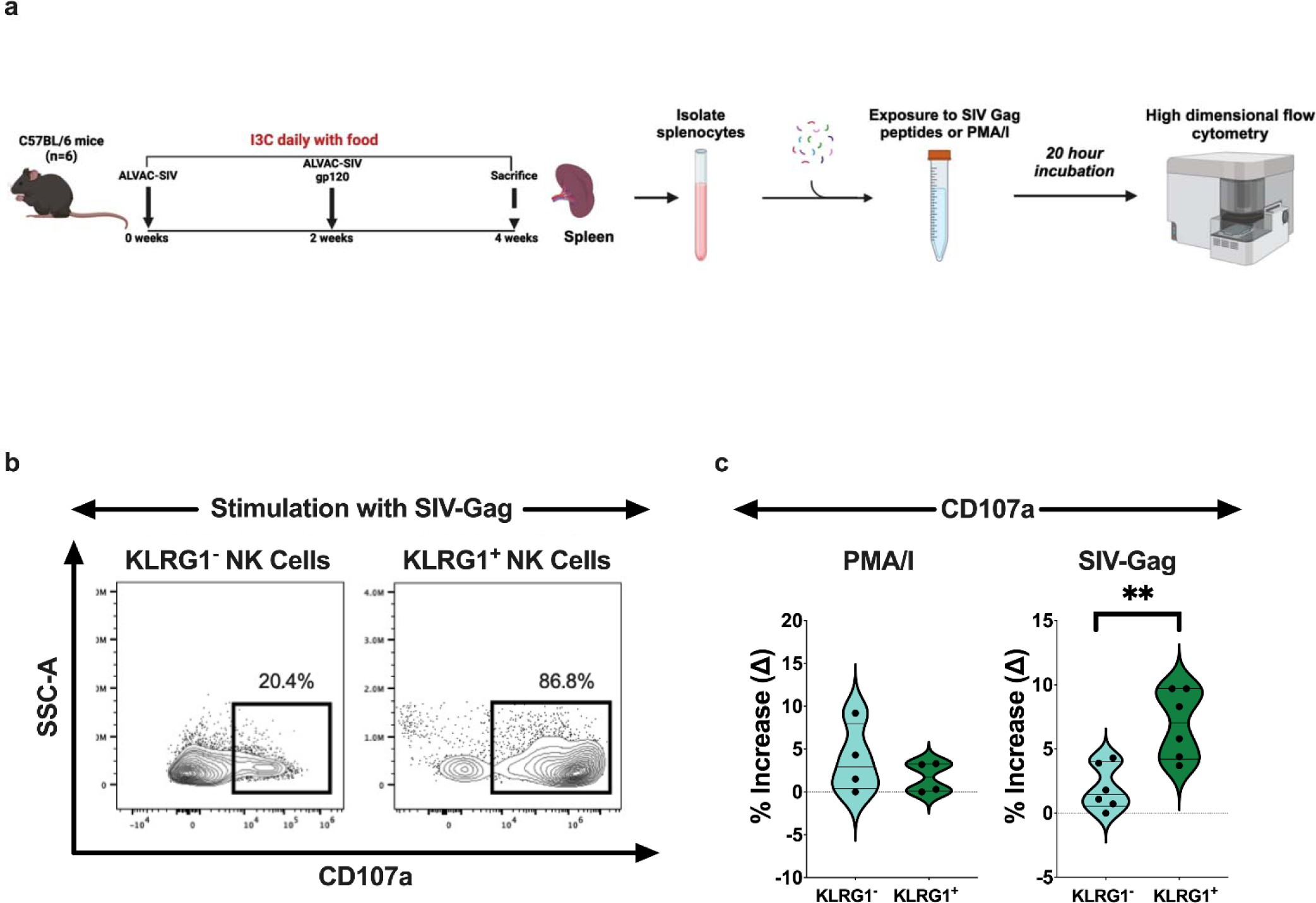
In vitro demonstration of recall responses of KLRG1^+^ (memory-like) NK cells to SIV peptides in mice administered an RV144-like vaccine with oral adjuvant I3C. **(a)** Experimental design. **(b)** Representative contour plots depicting CD107a expression in KLRG1^-^ and KLRG1^+^ NK cell subsets upon stimulation with SIV-Gag. **(c)** Violin plots depicting increase in expression (Δ values) of CD107a by KLRG1^-^ and KLRG1^+^ NK subsets upon stimulation with PMA/I and SIV-Gag peptides. The p-value was calculated using Wilcoxon rank-sum test; **p<0.01; PMA/I, phorbol 12-myristate 13-acetate (PMA) and ionomycin.

### Uncompromised Humoral Immune Responses Following Concurrent I3C Oral Adjuvant Administration

Given the current focus on broadly neutralizing antibodies in protection against HIV acquisition, well-functioning humoral responses are a critical component of HIV vaccines ^26^. These are also important in NK cell effector mechanisms such as ADCC, which has emerged as a correlate of protection in HIV vaccine responses. However, compelling evidence indicates that NK cells also impact B cell affinity maturation and immune function. Studies have shown that NK cell subsets may impair the humoral immune response by impeding the expansion and function of germinal center (GC) B cells ^27–30^. Given that our findings revealed I3C-mediated expansion of KLRG1^+^ NK cells with potentially superior effector functions, we sought to qualitatively assess the potential impact of I3C administration on the B cell response in the spleen of mice administered with an RV144-like vaccine regimen. We investigated selected B cell phenotypes which could be used to compare the functional competency of their vaccine induced humoral responses in the presence and absence of I3C. Here, we identified no significant difference in the frequency of total B cells between the two groups (Figure 4a). Interestingly we found several modulations in B cell surface receptor expression in the Vaccine + I3C mice which could be advantageous for a robust humoral response. These include significantly reduced PD-1 expression (p<0.05), indicating less inhibition of the B cell activation cascade ^31^(Figure 4b), significantly higher CD62L expression (p<0.01), suggesting a higher proportion of memory B cell subsets, likely against vaccine antigens ^32^ (Figure 4c), and greater Ly6C expression (p<0.05), suggesting a higher proportion of antibody secreting plasma cells within the B cell repertoire ^33^ (Figure 4d). However, we did not see any significant differences in B cell surface expression of CD27 and CD11c, which are markers for B cells’ expansion capability and potential for differentiation to plasma cells, respectively (Figures 4e-f) ^34,35^. Furthermore, using conventional flow cytometry gating, we identified subsets of KLRG1^-^ and KLRG1^+^ NK cells that are more likely to interact with germinal center responses based on CXCR5 and PD-1 expression (Figure 4g) ^36^. Our analysis revealed no significant difference in the frequencies of KLRG1^-^CXCR5^+^PD-1^+^and KLRG1^+^CXCR5^+^PD-1^+^ NK cells in the Vaccine + I3C group (Figure 4h), suggesting no greater potential of expanded KLRG1^+^ NK cells in interacting with germinal center B cells than their KLRG1^-^ counterparts. These findings collectively suggest that use of I3C as an oral adjuvant with vaccination is likely to maintain robust antibody responses required for a potentially effective HIV vaccine.

**Figure 4.**
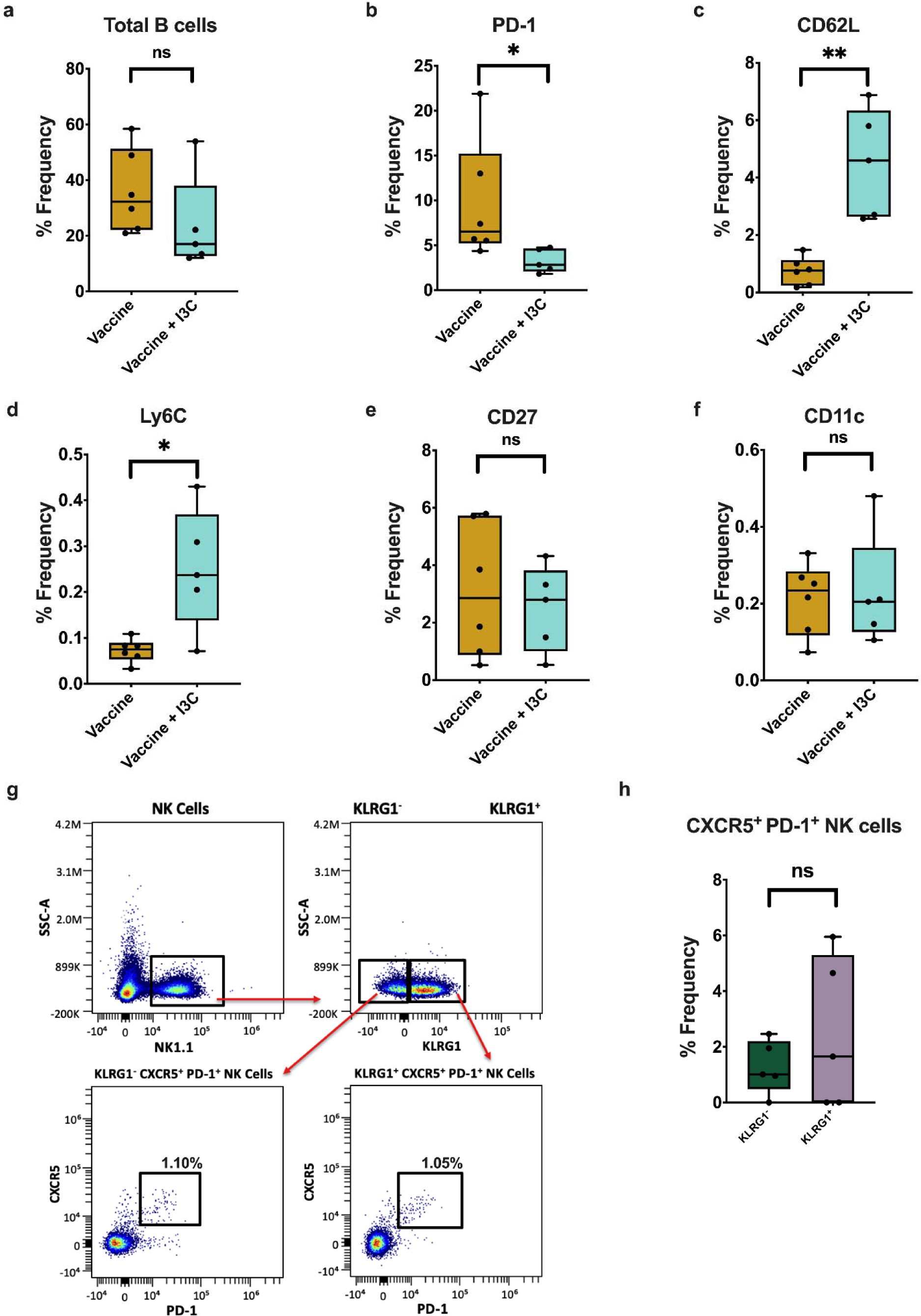
Uncompromised B cell responses when I3C used as an adjuvant with the RV144-like vaccine regimen. Box plots comparing **(a)** total B cells, and **(b)** PD-1^+^, **(c)** CD62L^+^, **(d)** Ly6C^+^**, (e)** CD27^+^, and **(f)** CD11c^+^ B cells (in total splenic NK cells) between Vaccine and Vaccine + I3C administered mice. **(g)** Representative pseudo color plots depicting the gating strategy for CXCR5^+^PD-1^+^ phenotypes in KLRG1^-^ and KLRG1^+^ NK cells. **(h)** Box plot comparing proportions of CXCR5^+^PD-1^+^ NK phenotypes among KLRG1^-^ and KLRG1^+^ NK cells in Vaccine + I3C administered mice. The p-value was calculated using Wilcoxon rank-sum test; *p<0.05, **p<0.01.

### Comprehensive Transcriptome Profiling Reveals Enhanced Antiviral Capabilities and ADCC Capacity of Splenic Immune Cells in mice That Received the Oral Adjuvant I3C Together with the Vaccine

We performed bulk RNA sequencing of splenocytes from both Vaccine and Vaccine + I3C groups. Our in-depth transcriptome analysis revealed 266 differentially expressed genes (DEGs) between the two groups (p<0.05) (Figure 5a). Among these DEGs, 187 were upregulated, and 79 were downregulated in the Vaccine + I3C group. Specifically, we observed upregulation of *Fcgr3* in the Vaccine + I3C group (Figure 5b), which encodes for FcγRIII, a receptor commonly found in NK cells and a critical component in the antibody-dependent cellular cytotoxicity complex ^37^. Similarly upregulated genes in this group include *Tnf* and *Ifngr2,* which encode a multifunctional proinflammatory cytokine belonging to the tumor necrosis factor (TNF) superfamily and a component of the IFNγ receptor, respectively. While TNFα is required for the lytic mechanism involved in ADCC, robust ADCC responses require efficient IFNγ-mediated responses ^38,39^. In the vaccine group administered I3C, we also observed upregulation of *Aim2* which encodes a cytoplasmic sensor of dsDNA, the activation of which may trigger pyroptosis in HIV infected cells containing viral dsDNA prior to integration ^40^. I3C also upregulated the gene encoding interferon-induced transmembrane proteins 2 (IFITM2) that potently inhibits HIV-1 replication by interfering with viral entry ^41^. Other upregulated genes included those encoding certain chemokine ligands such as *Ccl6*, which promote innate immune cell activation and recruitment ^42^. Collectively, these suggest that the incorporation of I3C to our RV144-like vaccine regimen potentially enhances the ability of the tissue environment to resist HIV/SIV acquisition. Further, we noted an upregulation of genes encoding structural constituents of chromatin, such as *Hist1h2ac,* which may reflect an alteration in the complexity in chromatin structure and function, suggesting that certain transcriptional changes induced by concurrent administration of I3C may result from epigenetic reprogramming in splenocytes, predicting the potential for long-term persistence of these changes, possibly in the form of innate immune memory ^43,44^.

**Figure 5.**
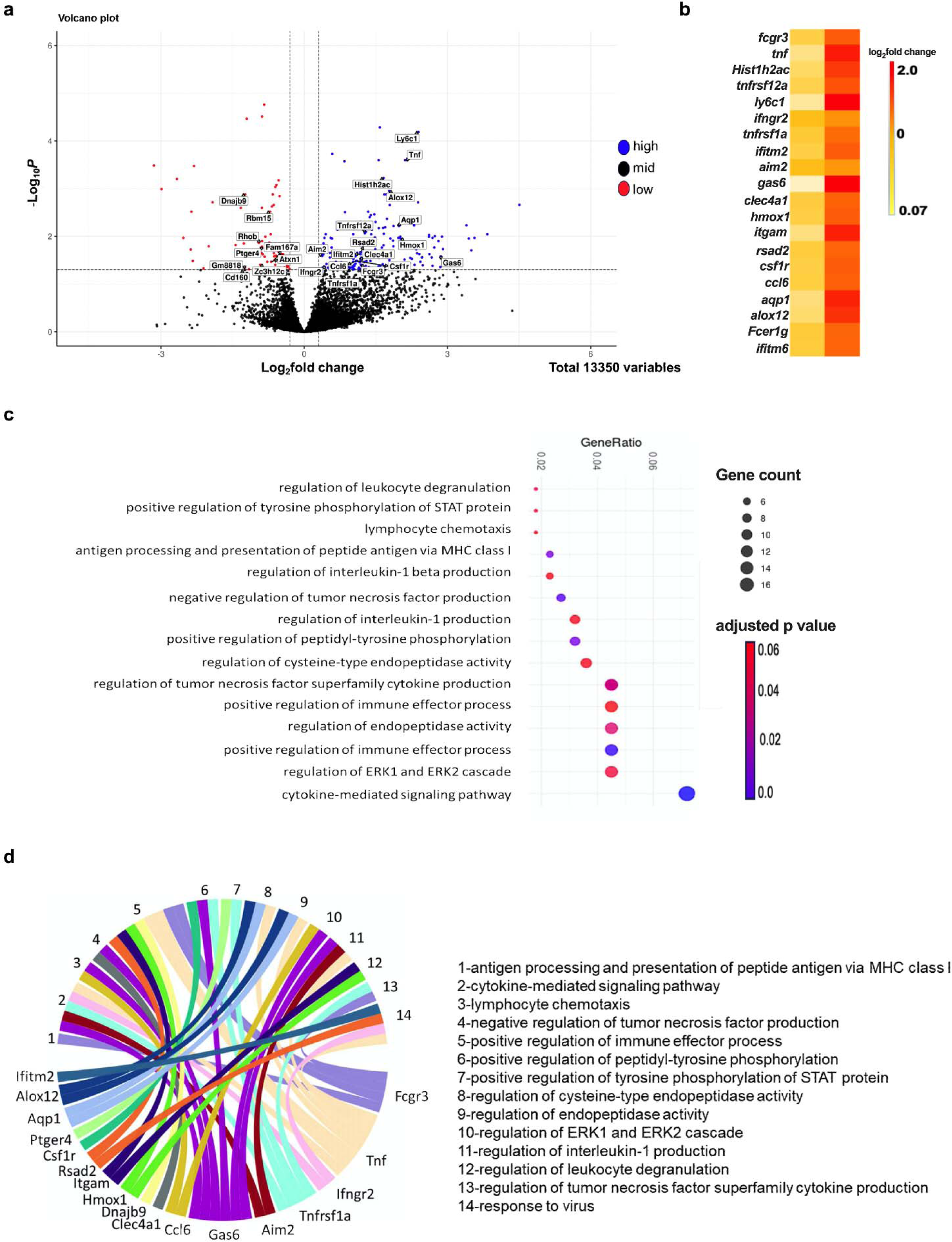
Transcriptome analysis reveals enhanced antiviral capabilities and capacity for ADCC when I3C used as an oral adjuvant. **(a)** Volcano plot depicting differentially expressed genes (DEGs) of splenocytes between Vaccine and Vaccine + I3C administered mice. **(b)** Heatmap highlighting upregulated DEGs in the Vaccine + I3C that may promote the ADCC capacity of NK cells. **(c)** Kyoto Encyclopedia of Genes and Genomes (KEGG) pathway analysis generated using DEGs to identify pathways of biological processes that were enriched in the Vaccine + I3C group, compared with the Vaccine group. **(d)** Circos plots exploring relationships between DEGs and enriched pathways of biological processes among the analyzed groups.

To examine the alterations in transcriptional pathways of biological processes influenced by these DEGs, we performed a Kyoto Encyclopedia of Genes and Genomes (KEGG) pathway analysis (Figure 5c). We found several pathways related to antiviral and cytotoxic functions of immune cells to be differentially enriched between the Vaccine and Vaccine + I3C groups. These included leukocyte degranulation, lymphocyte chemotaxis, immune effector process regulation, cytokine-mediated signaling pathways, regulation of TNF superfamily cytokine production, regulation of IL-1β production, antigen processing and presentation, regulation of tyrosine phosphorylation of STAT protein, and ERK1 and ERK2 cascade regulation (p<0.05 for all). Enrichment of these pathways indicate that the administration of oral adjuvant I3C in concomitance with vaccination has a significant impact on antiviral and cytotoxic immune cell effector processes. The intricate connections observed between several upregulated genes, as a result of concurrent I3C administration, and enriched pathways regulating immune cell antiviral responses and cytotoxicity (Figure 5d) offers insights into the impact of I3C in manipulating immune cell effector responses against microbial pathogens, including HIV, at a transcriptional level.

### I3C May induce phenotypic and functional enhancement in KLRG1^+^ NK through Epigenetic Modifications

We observed the expansion of a subset of KLRG1^+^NK cells with enhanced vaccine antigen-specific memory-like features, coupled with transcriptomic evidence of chromatin remodeling of splenocytes during concurrent I3C administration with the vaccine. These unique observations suggest the possibility of underlying epigenetic modifications of NK cells induced by the AHR agonist I3C (Figure 6a). Due to lack of cell number, we were unable to explore this further through our mouse model. Nevertheless, to evaluate if I3C directly regulates NK cells to induce the expression of unique phenotypes, we examined publicly available AHR agonist ChIP-seq in lymphocytes, as well as our previously published chromatin assay for transposase-accessible chromatin with sequencing (ATAC-Seq) done on human NK cells ^45,46^. We found that AHR agonists do bind to the KLRG1 promoter, which overlaps the site of chromatin accessibility in human NK cells (Figure 6b). Thus, it remains possible that I3C could be directly acting on NK cell epigenetic programs.

**Figure 6.**
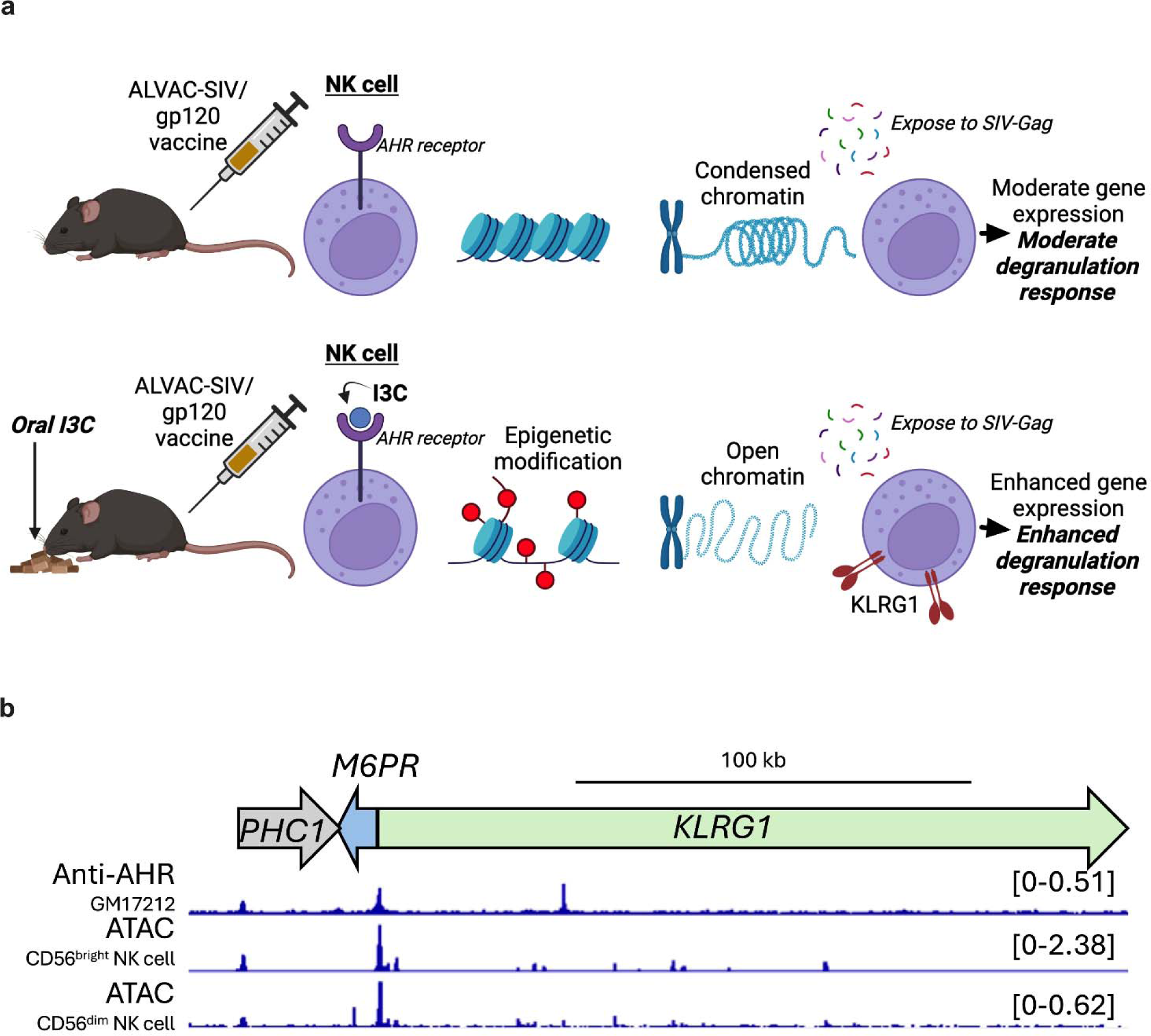
AHR agonists (I3) may induce phenotypic and functional enhancement in KLRG1^+^NK cells through epigenetic modifications. **(a)** Schematic diagram representing hypothesis for phenotypic and functional enhancement of vaccine antigen specific KLRG1^+^ NK cells by concurrent oral I3C administration to mice. **(b)** Integrative Genomics Viewer (IGV) snapshot showing the human KLRG1 gene region. The scale and relative gene locations are shown on top. The histograms represent anti-AHR ChIP-seq data, derived from a lymphoblastoid cell line termed GM17212, and assay for transposase-accessible chromatin with sequencing (ATAC-Seq) data from human CD56^bright^ NK cells or CD56^dim^ NK cells.

## Discussion

In this study, using a mouse model, we found that use of I3C as an oral adjuvant with a RV144-like vaccine regimen resulted in the expansion of KLRG1^+^ NK cells with phenotypic characteristics reflecting an overall greater cytotoxic potential (CD11b^hi^CD27^low^) compared to their KLRG1^-^ counterparts. Subsequent in vitro experiments demonstrated that the expanded KLRG1^+^ NK subset displayed a comparatively enhanced cytotoxic potential upon stimulation with HIV peptides, suggesting a memory-like responses to vaccine antigens. Transcriptome analysis further revealed that the immune system displayed an overall greater capacity to prevent HIV acquisition, through multiple antiviral mechanisms, including those such as ADCC, when I3C was used as an oral adjuvant with the RV144-like vaccine regimen. This was marked by upregulation of genes associated with several immune effector processes.

Previous studies have identified KLRG1 as a biomarker of NK cell activation ^20^, with subsets exhibiting memory-like features associated with protection against infectious organisms like *Mycobacterium tuberculosis* ^47^. Jost et. al recently demonstrated increased KLRG1 expression in HIV-specific NK cell subsets among HIV-infected individuals indicating that KLRG1 can be considered a useful biomarker of HIV/SIV-specific memory NK cells ^48^.

We further delineate the phenotypic and functional characteristics of these expanded KLRG1^+^ NK cells induced by I3C in comparison to KLRG1^-^ NK cells. Here, we found KLRG1^+^ NK cells to display an overall enhanced cytotoxic potential, as evidenced by higher expression of CD11b and CD11c and lower CD27 expression ^16,21^. Their comparatively higher expression of Ly6C on the other hand indicates that these KLRG1^+^ NK cells have high plasticity and represent a reservoir of effector NK cells which has the potential to produce an effective and strong response to infection ^23^.

Moreover, our in vitro experiments revealed a greater increase in the degranulation response in Vaccine + I3C-exposed KLRG1^+^ NK cells compared to their KLRG1^-^ counterparts, specifically upon stimulation with vaccine antigens, while no significant difference was observed upon non-specific stimulation. This suggests these cells are preferentially activated as a result of antigenic memory. Moreover, our transcriptomics findings provide evidence for chromatin remodeling of splenocytes upon addition of I3C to the vaccine regimen. This, along with independent evidence of AHR agonists impacting KLRG1 promoter regions suggest that KLRG1 expression may denote a subset of splenic NK cells potentially undergoing epigenetic modifications induced by the vaccine in combination with I3C. However, to further ascertain this, investigation of this subset of KLRG1^+^ NK cells exposed to both the vaccine and oral adjuvant I3C, particularly through ATAC-Seq analysis, is required. This would identify unique epigenetic modifications which may have occurred in this subset (compared to KLRG1^-^ NK cells) as a result of I3C being used as an oral adjuvant with the vaccine regimen indicating long-term epigenetic reprogramming of these cells (i.e., trained immunity) ^49^.

Considering that NK cells are the key effector cells in antiviral responses though mechanisms such as ADCC, robust antiviral responses are improved by competent NK cells with enhanced cytotoxicity. Previous literature suggests that KLRG1^+^ NK cells exhibit a heightened capacity to mediate antiviral mechanisms such as ADCC ^15^. Moreover, their enhanced cytotoxic nature, particularly in response to vaccine antigen-induced recall responses, implies that expanded KLRG1^+^ NK cells may excel in such antiviral effector mechanisms against HIV/SIV infected cells^50^. This was supported by our transcriptomic analysis which revealed an environment conducive for efficient ADCC (upregulated *Fcgr3* and *Tnf*) and other antiviral effects (upregulated *Aim2*, *Ifitm2*, *Ccl6*), and enrichment of pathways related to regulation of leukocyte degranulation and immune effector processes following I3C used as an oral adjuvant with the HIV/SIV vaccine. Additionally, while studies have demonstrated that the ALVAC vector itself used in our study primes *Aim2* triggering strong inflammasome activation, I3C seems to bolster the vaccine’s inherent antiviral effects through *Aim2* upregulation, further justifying the use of I3C as an adjuvant to an ALVAC vector-based vaccine strategy ^51^.

Collectively, our findings suggest that, orally administered I3C promotes the expansion of KLRG1^+^ NK cells, a subset of vaccine antigen specific memory-like NK cells with an enhanced cytotoxic capacity. This enrichment does not seem to compromise humoral responses but augments the potential to effectively mediate NK cell mediated anti-HIV/SIV effector mechanisms. Being already identified as a correlate of protection in the RV144 vaccine regimen, this greater potential for robust effector mechanisms like ADCC in expanded vaccine antigen specific memory-like NK cells is likely to increase and sustain the efficacy of HIV vaccine regimens.

Our study has several limitations. First, our comprehensive immune analysis was restricted to splenic NK cells due to inadequate cell numbers for such an analysis in other tissues. Second, we were unable to specifically demonstrate the enhancement of NK cell effector mechanisms like ADCC and the killing ability of HIV/SIV-infected cells by KLRG1^+^ NK cells from the Vaccine + I3C group through mechanistic studies due to limitations associated with a mouse model. Third, our transcriptome analysis was based on bulk RNA sequencing data and thus we cannot make cell-specific extrapolations from this. However, our findings of I3C-induced expansion of a subset of NK cells with vaccine antigen-specific cytotoxic memory and its potentially broad impact on enhancing antiviral immunity in systemic tissues (as evidenced by our transcriptome analysis) warrant further investigation into the potential of I3C and similar compounds as adjuvants to enhance the efficacy of existing HIV vaccines. Further studies using non-human primate-SIV/HIV models are required to validate these findings.

## Methods

### Ethics Statement

This study and its associated experiments were approved by The Ohio State University’s (OSU’s) Institutional Biosafety Committee (IBC) and Institutional Animal Care and Use Committee (IACUC). The study performed aligned with relevant institutional guidelines and is reported in accordance with ARRIVE guidelines (https://arriveguidelines.org)

### Mouse Strains

CD57BL/6 mice, aged 4-6 weeks (six males and six females), were obtained from The Jackson Laboratory (Bar Harbor, ME). All mice were maintained within OSU’s Laboratory Animal Resources (ULAR) and supplied with sterile and water ad libitum. All methods and procedures performed were in accordance with OSU’s Institutional Animals Care and Use Committee’s (IACUC’s) approved protocols and procedures.

### Immunization

A vaccine regimen similar to that utilized in the RV144 clinical trial was administered to all mice^3^. The vaccine administered intramuscularly, utilized the ALVAC (vCP2432) expressing SIV genes *gag-pro* and *gp120TM* (Lot No: 032812-4), and a bivalent monomeric-gp120 protein boost consisting of SIV_mac251_-M766 gp120-gD formulated in alum (Lot no. 102717A).

### Experimental Design

Mice were evenly segregated into two groups (3 males and 3 females in each) and fed a rodent diet (AIN-76A) throughout the study period. All mice received the following vaccine regimen, intramuscularly to the thigh: ALVAC-SIV (10^8^ PFU in 50μL) at week 0 and week 2, and gp120 (50 μL) in alum (to the opposite thigh) at week 2. The second group (group B) (Vaccine + I3C) additionally received daily oral administration of 100 ppm I3C, with food for 4 weeks starting at week 0.

### Tissue Collection and Processing for Flow Cytometry

Mice were humanely euthanized at week 4 and their spleens were collected for cell isolation as described previously^52^. Spleens were gently sieved through a mesh screen using the plunger portion of a 1 mL syringe. The resultant cell suspensions were then transferred to a 15 mL tube by filtering through a 100-micron filter using ice-cold R10 media (RPMI 1640 by Gibco® Thermo Fisher, 10% FBS, 2 mM l-glutamine, 100 U/mL penicillin G, 100 μg/mL streptomycin). Followed by a spin-down of the tubes at 250×g for 7 min at 4°C. The cell pellet was next resuspended with R10 media. After discarding the supernatant, the cell pellet was once again washed and incubated with 1 mL of RBC lysis buffer at room temperature (All washing procedures were performed at 700xg for 5 min at 4°C). The RBC lysis reaction was stopped after 10 minutes using 1 mL of R10 media and spun down at 250×g for 7 min at 4°C.

Freshly isolated splenocytes were transferred to pre-warmed R10 media and washed. Cells were then resuspended at 1–2 million cells per ml in R10 media and was followed by surface staining for flowcytometry as follows: Samples were stained with Fixability Viability Dye (Zombie NIR Cat No. 423105) and incubated for 15 minutes. Cells were then washed, followed by the addition of a surface antibody cocktail (Supplementary Table 1). Antibodies utilized were previously titrated to determine the optimal concentration. After a 30-minute incubation period, cells were washed again. Following the filtering of cells through strainer capped FACS tubes, samples were acquired on a Cytek Aurora flow cytometer. Monoclonal antibodies used in the panel can be accessed in Supplementary Table 1.

### Flow Cytometry Data Analysis

FCS files were exported from SpectralfFo and imported into Omiq for subsequent analysis. We obtained “Fluorescence Minus 1” controls (cells stained with all fluorochromes used in the experiment except 1) for each marker prior to spectral unmixing (Supplementary figure 2b).

### In Vitro Assay for NK Cell Degranulation

Splenocytes from Vaccine + I3C mice were cultured in RPMI-1640 medium supplemented with 10% fetal bovine serum (FBS), 100 U/mL penicillin, and 100 µg/mL streptomycin and incubated at 5% CO_2_ at 37°C. Cells from each sample were divided equally to two aliquots and exposed to either SIV-gag peptides (10µg) or phorbol 12-myristate 13-acetate (PMA) and Ionomycin (eBioscience^TM^, Cat. No. 00-4970-93 1µL/mL) and incubated for 20 hours. Post-incubation, cells were harvested from the culture wells and stained with fluorochrome-conjugated antibodies against CD107a. Flow cytometry was performed to quantify the levels of CD107a expression on different NK cell subsets.

### RNA Extraction and Sequencing

The cell pellets were transported to Genewiz/Azenta in South Plainfield, New Jersey, for processing. Library preparation was performed using the Illumina sequencing platform. The chosen method was Standard RNA-sequencing with PolyA selection, aiming for a sequencing depth of 20 million reads per sample. Subsequently, the analysis encompassed trimming of sequence reads, mapping of the trimmed reads and evaluation of differential gene expression to identify genes that exhibited statistically significant changes in expression across various conditions or treatments. Following these initial analyses, further investigations were conducted using R Studio version (4.2.1), employing packages such as ggplot, enrichGO, and ggpubr to facilitate visualization and enrichment analysis of the gene expression data.

### ChIP-seq and ATAC-seq analysis

AHR agonist ChIP-seq (GEO ID: GSM3244480) and NK Cell ATAC-seq (GEO IDs: GSM3484011 and GSM3084217) data have been published previously (37,38). Data were downloaded from the GEO website, aligned to the Hg38 genome using Bowtie2 (v 2.2.9), and converted into counts per million visualizations using deepTools (v 3.5.6). The resulting tracks were displayed on the IGV genome browser (v2.16.1) and then manually annotated using Microsoft PowerPoint.

### Graphing and Statistics

Graphs were prepared using Graph Pad Prism (version 10). Data was statistically analyzed using their bundled software. Statistical comparisons were performed using the Wilcoxon rank-sum test. Differences between groups were considered significant if p<0.05 (graphically indicated by one or more asterisks; one asterisk represents p<0.05 and two asterisks represents p <0.01). All t-tests were two-tailed and nonparametric tests were used with a confidence interval of 95%. Each in vivo analysis was performed with six mice per group as determined by a power calculation using the assumption (based on prior data) that there will be at least a twofold change with a standard deviation of <0.5. Error bars represent the meanL±Lthe standard error of the mean.

## Data Availability

The data sets generated and analyzed during the current study have been uploaded to GitHub and are accessible through Zenodo under the DOI: 10.5281/zenodo.11087021.

## Acknowledgments

We would like to express our sincere gratitude to Dr. Genoveffa Franchini NIH/NCI for generously providing the ALVAC-SIV vaccine and gp120 protein for this study. This study was supported by NIAID/K22 AI127072 award to N.L and The Ohio State University, Infectious Diseases Institute Host Defense and Microbial Biology Pilot Grant award to N.L.

## Author Contribution

M.G, M.A, and C.I, performed experiments, data analysis and wrote the manuscript. I.D. performed data analysis and wrote the manuscript. S.B and W.M. performed experiments, data analysis, helped with literature search and manuscript writing. T.D designed the study and edited the manuscript. P.C. performed epigenetic analysis and edited the manuscript. N.L. supervised the project, designed the study, performed data analysis, and wrote the manuscript.

