## Supplemental Figures for "Novel Oral Adjuvant to Enhance Cytotoxic Memory-Like NK Cell Responses in an HIV Vaccine Platform"

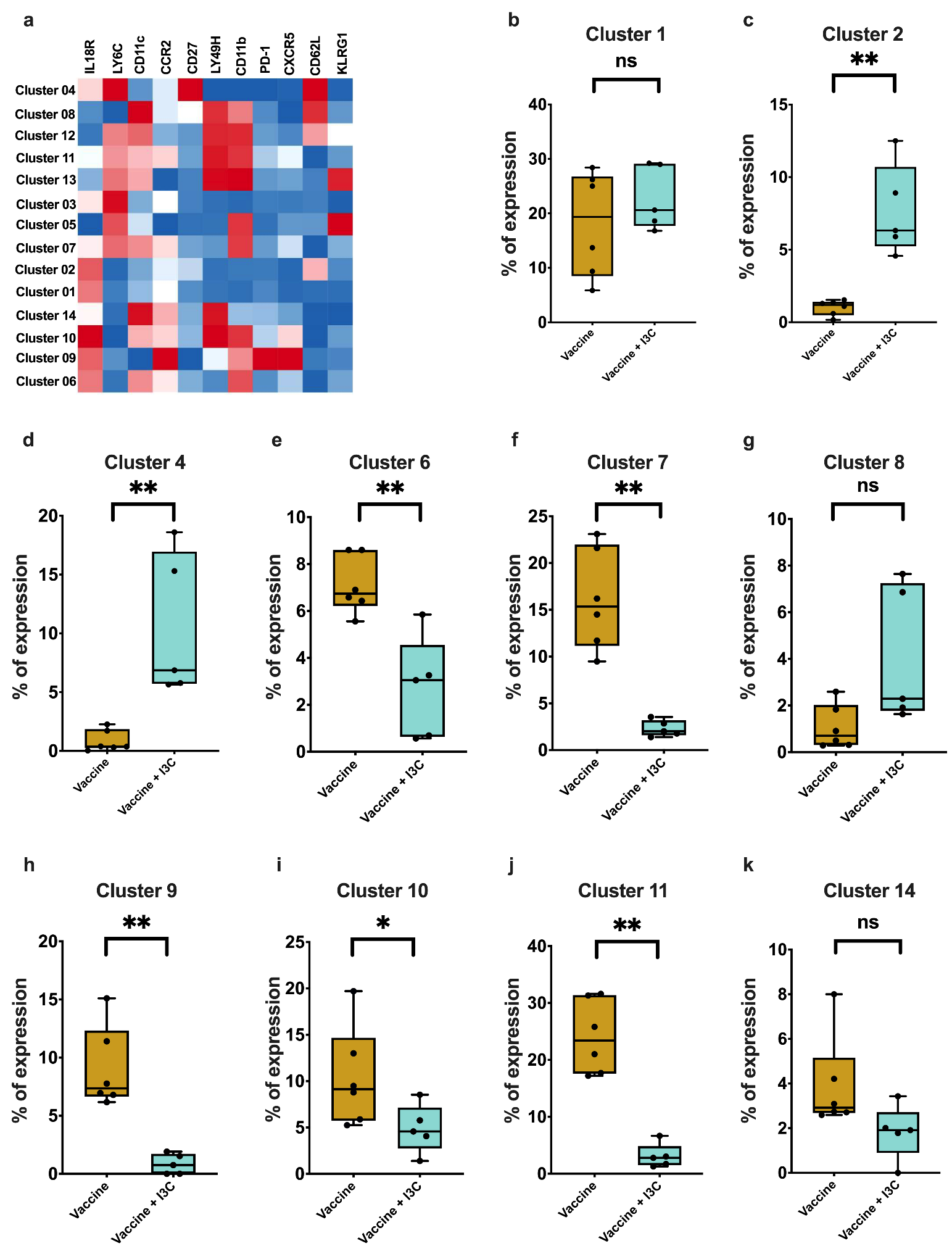

**Supplementary figure 1. (a)** Clusters-by-marker heatmap characterizing the receptor expression patterns of individual NK cell clusters. **(b-k)** Box plots depicting FlowSOM-based NK cell cluster frequency differences between Vaccine and Vaccine + oral adjuvant I3C groups. The p-value was calculated using Wilcoxon rank-sum test; *p<0.05, **p<0.01. I3C, Indole-3-carbinol.

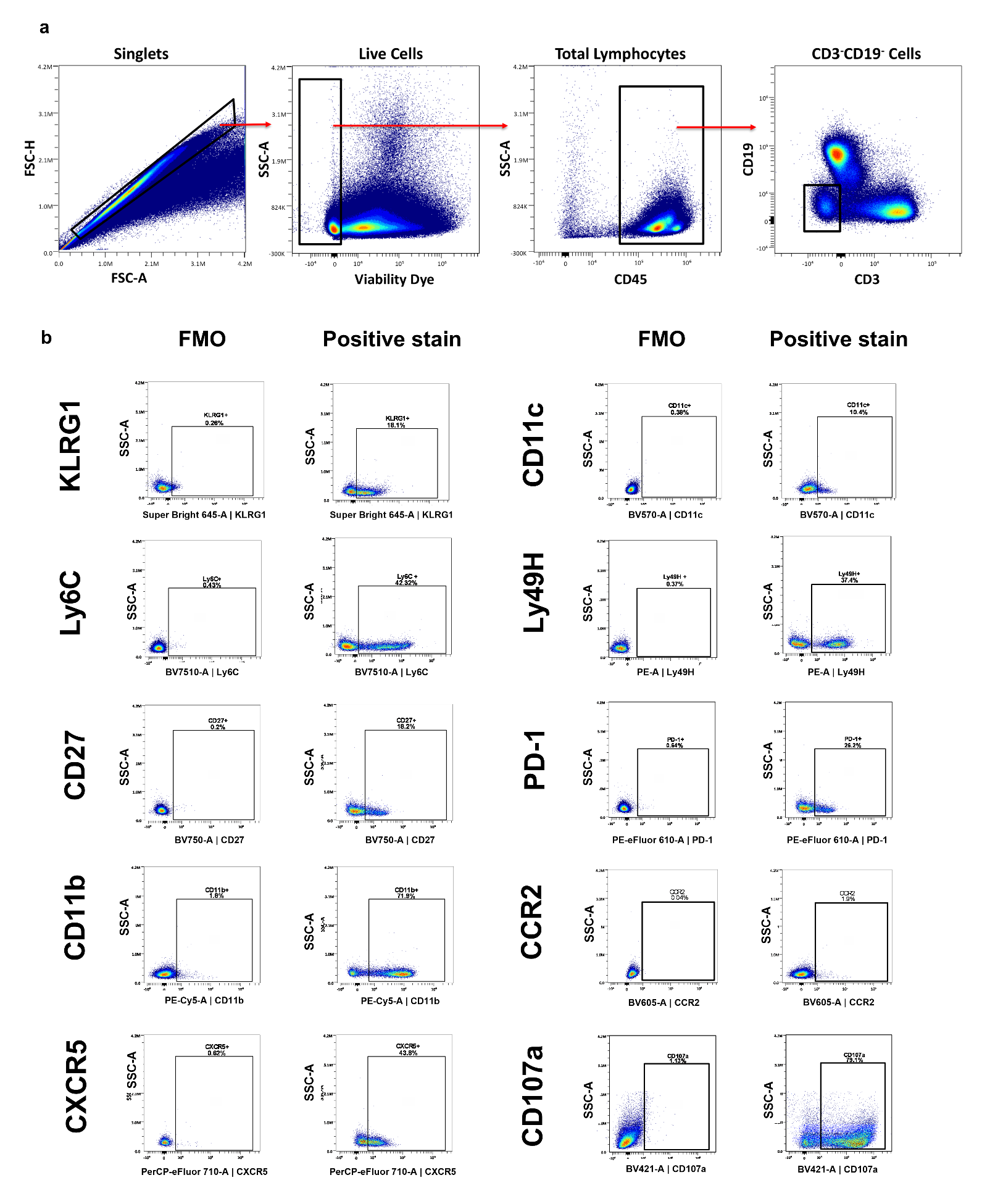

**Supplementary figure 2. (a)** Representative pseudo color plots depicting flow cytometric gating strategy used to define immune cell populations in mouse splenocytes. **(b)** Representative pseudo color plots illustrating fluorescent minus one control for receptor expression.
