## Supplemental Tables for "Novel Oral Adjuvant to Enhance Cytotoxic Memory-Like NK Cell Responses in an HIV Vaccine Platform"

| **Fluorochrome** | **Specificity** | **Clone** | **Company** | **Catalog No.** | **Concentration** |
| --- | --- | --- | --- | --- | --- |
| FITC | CD3 | 17A2 | Biolegend | 100203 | 1:50 |
| PE- eFluor 610 | PD-1 | J43 | Invitrogen | 61-9985-82 | 1:50 |
| PE-Cy5 | CD11b | M1/70 | Invitrogen | 15-0112-83 | 1:50 |
| PerCP-eFluor 710 | CXCR5 | SPRCL5 | Invitrogen | 46-7185-82 | 1:50 |
| Brilliant Violet 421 | CD45 | 30-F11 | Biolegend | 103134 | 1:50 |
| Brillant Violet 605 | CCR2 | SA203G11 | Biolegend | 150615 | 1:50 |
| Super Bright 645 | KLRG1 | 2F1 | Invitrogen | 64-5893-82 | 1:50 |
| Super Bright 436 | CD62L | MEL-14 | Invitrogen | 62-0621-82 | 1:50 |
| Brilliant Violet 711 | CD19 | 6D5 | Biolegend | 115555 | 1:50 |
| Brilliant Violet 750 | CD27 | LG.3A10 | BD | 747399 | 1:50 |
| Brilliant Violet 785 | NK1.1 | PK136 | Biolegend | 108749 | 1:50 |
| Brilliant Violet 510 | Ly6C | HK1.4 | Biolegend | 128033 | 1:50 |
| Brilliant Violet 570 | CD11c | N418 | Biolegend | 117331 | 1:50 |
| PE | Ly49H | 3D10 | Biolegend | 144705 | 1:50 |

**Supplementary Table 1:** Monoclonal Antibodies used for flow cytometry
